# Climate change impacts the structure and nitrogen-fixing activities of subarctic feather moss microbiomes across a precipitation gradient

**DOI:** 10.1101/2025.06.06.658193

**Authors:** Danillo O. Alvarenga, Justin T. Wynns, Joseph Nesme, Anders Priemé, Kathrin Rousk

## Abstract

Associations between feather mosses and cyanobacteria are crucial sources of new biologically available nitrogen (N) in arctic and subarctic ecosystems. The physiology of both mosses and cyanobacteria is strongly influenced by environmental factors such as temperature and moisture, which directly affect N_2_ fixation rates. These associations may be threatened by climate change, since it leads to warmer and drier conditions in polar regions. In this study, we investigated the N_2_-fixing microbial communities associated with two common feather mosses across a precipitation gradient in the subarctic tundra, followed by a temperature and moisture experiment. Using acetylene reduction assays, *nifH* gene sequencing and qPCR, we evaluated how shifts in temperature and moisture influence nitrogenase activity and N_2_-fixing community structure. Our results showed that N_2_ fixation was highest in sites with greater precipitation and increased with both temperature and moisture. Cyanobacteria dominated N_2_-fixing communities, but currently unclassified bacteria also seemed to play a significant role, particularly at high temperatures. The number of cyanobacterial *nifH* copies tended to decrease with temperature, while the relative abundance of unclassified bacteria increased. These findings suggest that the N_2_-fixing activity, abundance and diversity of cyanobacteria associated with feather mosses in the subarctic will decline under warmer and drier conditions, potentially leading to a shift in the composition of feather moss-associated microbial communities in a warmer Arctic, with potential consequences for N input into the ecosystem.

## INTRODUCTION

Mosses play a crucial ecological role in environments characterized by cold temperatures and low abundance of vascular plants, such as tundra ecosystems. In the arctic and subarctic tundra, perennial moss species often dominate the ground cover, driving carbon sequestration and hosting nitrogen-fixing cyanobacteria (Solheim et al., 2004; Douma et al., 2007; Street et al., 2013; Eldridge et al., 2023). Associations between mosses and cyanobacteria introduce significant amounts of new N into tundra ecosystems despite low temperatures, high rates of ultraviolet radiation during the growing season and fluctuating water availability (Kvíderová et al., 2019). However, the contributions of mosses and their associated cyanobacteria to C and N cycles in these environments are increasingly threatened by climate change.

Arctic regions are warming much faster than the rest of the planet due to the polar amplification phenomenon, largely resulting from the loss of sea ice (Dai et al., 2019; Rantanen et al., 2022; Xie et al., 2022), and currently experience their highest temperatures on record (Ballinger et al., 2023). One of the most alarming consequences of the increasing temperatures in these regions is lower humidity and increased frequency of drought events (Finger Higgens et al., 2019). Both moss and cyanobacteria are poikilohydric organisms, and as such they are incapable of regulating their moisture levels, depending entirely on environmental conditions to stay hydrated. Consequently, C sequestration by mosses is highly vulnerable to drought, which decreases their net primary production (Martínez-García et al., 2024). The efficiency of N_2_ fixation in moss-cyanobacteria associations is also significantly impacted by abiotic factors, especially temperature and moisture (Rousk et al., 2017a; Rousk, 2022). Nevertheless, the effects of climate change on moss-cyanobacteria associations in arctic and subarctic ecosystems are still unclear.

Some works have shown that climate change directly and indirectly reduces N_2_ fixation, cyanobacterial colonization and ground cover associated with feather mosses in subarctic ecosystems (Sorensen & Michelsen, 2011; Sorensen et al., 2012; Alvarenga & Rousk, 2021; Permin et al., 2022). On the other hand, other studies estimated that some of the effects of climate change, like higher temperatures and drier habitats, may actually increase N_2_ fixation associated with subarctic mosses (Lett & Michelsen, 2014; Rousk & Michelsen, 2017; Rousk et al., 2017a). In addition, mosses may buffer some ecosystems against the consequences of climate change by storing water, C and N (Slate et al., 2024). These contrasting results indicate that the effects of climate change on the composition and productivity of these N_2_-fixing associations are still poorly understood. This information is crucial for accurate predictions of how will N-limited ecosystems be impacted by changing environmental conditions.

Understanding how climate change will affect the contributions of moss-cyanobacteria associations to biogeochemical cycles is further complicated by the fact that mosses host distinct microbial communities. Moss microbiomes can vary based on both the identity or evolutionary history of the host species and environmental factors such as light or temperature (Holland-Moritz et al., 2021). Warming may also drive shifts in the composition of N_2_-fixing microbial communities (Klarenberg et al., 2022). Furthermore, moss microbiomes contain novel microbes with yet unclear ecological functions (Holland-Moritz et al., 2018), introducing an element of unpredictability into our understanding of how these communities may respond to climate change. This variability in microbial composition poses significant challenges for predicting how N_2_ fixation will respond to climate change across various moss species and environments.

In this study, we evaluated how climate change-driven temperature and water stress impacts N_2_ fixation associated with two dominant feather mosses from subarctic tundra. We sampled two feather moss species growing across a natural precipitation gradient in northern Sweden and analyzed N_2_ fixation rates, N_2_-fixing community structure and *nifH* gene abundance under three different temperatures. We hypothesized that: 1) N_2_ fixation rates are higher in feather mosses from sites with higher annual precipitation; 2) N_2_ fixation rates are positively correlated with higher temperatures and moisture levels; and 3) these potential increases in N_2_ fixation are positively correlated with the larger abundance of cyanobacteria in the feather moss microbiomes.

## METHODS

### Overview of sampling sites

This study targeted three different sites in northern Sweden, representing a steep precipitation gradient within 40 km: a low precipitation site close to Abisko (hereafter referred to as ABK) (68°20’57.7”N, 18°49’48.1”E); a medium precipitation site close to Låktajåkko (LTJ) (68°24’54.6”N, 18°24’21.9”E); and a high precipitation site close to Katterjåkk (KTJ) (68°25’10.1”N, 18°09’51.5”E). According to the Swedish Meteorological and Hydrological Institute (https://www.smhi.se/), sites ABK, LTJ and KTJ have mean annual precipitations of 300 mm, 800 mm and 1200 mm, respectively. Annual mean temperatures are 0.3 °C for ABK, −3.4 °C for LTJ and −0.7 °C for KTJ (https://www.smhi.se/). Given the proximity of the sites, other factors are similar, such as soil pH, which was around 6.0 (6.0±0.2 at ABK, 6.1±0.1 at LTJ and 5.9±0.2 at KTJ). The three sites were similar in vegetation, being often dominated by two feather moss species, *Hylocomium splendens* (Hedw.) Schimp. and *Pleurozium schreberi* (Brid.) Mitt., as well as the shrubs *Empetrum hermaphroditum* Hagerup, *Vaccinium uliginosum* L. and *Betula pubescens* Ehrh.

### Sample collection and incubation

Sampling was carried out in October 2016, just before the first snowfall in northern Sweden. Six 18×18×18 cm mesocosms from each site containing the two dominant feather moss species, *H. splendens* and *P. schreberi*, were collected and placed in white plastic boxes. *H. splendens* was not found at LTJ, therefore only *P. schreberi* was sampled. The mesocosm boxes were shipped to the University of Copenhagen for downstream experimental work. Randomly-selected 3 g portions of moss shoots from each mesocosm were placed in sterile 50 mL transparent plastic tubes and kept at different temperature regimes (5, 15 or 25 °C). As a full-factorial set-up, for each temperature, the samples were also exposed to the moisture levels of 50 (consisting of partially hydrated shoots), 100 (fully hydrated shoots) or 150 % (fully hydrated shoots with an extra volume of water corresponding to half of its moisture). Six replicates were used in the different treatments, resulting in 270 samples. The samples were kept in growth chambers under a photoperiod of 12 h of light at 200±25 µmol photons·m^−2^·s^−1^ and 12 h of darkness for 8 weeks, being monitored for water loss once every week.

### Acetylene reduction assays

We performed acetylene reduction assays (ARA) as an indirect measure of N_2_ fixation in the samples (Hardy & Knight, 1967). Each tube prepared in the previous step was sealed with a rubber Suba Seal septum (Sigma-Aldrich, Saint Louis, USA) and 10 % of the headspace was replaced with acetylene gas (i.e., 50 mL). The vials were incubated under the previous conditions for durations that varied with incubation temperatures: samples at 5 °C were incubated for 18.5 h; samples at 15 °C, for 7.5 h; and samples at 25 °C, for 3 h. Six mL of the vial headspaces were transferred into pre-evacuated, air-tight vials (Labco, Lampeter, UK) and analyzed for ethylene concentrations with an SRI 310C FID gas chromatograph (SRI Instruments, Torrance, USA). Gas chromatography was performed after the samples were incubated for 1 day (1D), 1 week (1W), 2 weeks (2W), 4 weeks (4W) and 8 weeks (8W).

### DNA isolation

After 8 weeks of incubation, the moss shoots were freeze-dried, resulting in an average sample weight of 30.2±0.1 mg. Five of the six ARA replicates in each treatment (225 samples) were selected for subsequent qPCR and DNA sequencing. The greenest parts of the feather moss shoots were separated from older portions and discolored tips, if present, and any organic material not belonging to the target moss species was removed. These samples were cut into fine fragments with sterile scissors and transferred to PowerBead Pro tubes (Qiagen, Hilden, Germany). The cells were broken in the FastPrep-24 benchtop homogenizer (MP Bio, Solon, USA) at a rate of 5.5 m·s^−1^ for 40 s and DNA isolation was carried out with the DNeasy PowerSoil Pro Kit (Qiagen) following manufacturer’s instructions. Isolated DNA was screened on 1 % agarose gels and stored at –20 °C.

### *nifH* gene amplification and sequencing

Amplification of the *nifH* gene in the isolated DNA was performed using a inosine-free modification of the primers 19F (5’-GCN WTY TAY GGN AAR GGN GG-3’) and 407R (5’-AAN CCR CCR CAN ACN ACR TC-3’) by Ueda et al. (1995). These primers were further adapted for indexing with upstream additions of Illumina overhang adapter sequences (5’-TCG TCG GCA GCG TCA GAT GTG TAT AAG AGA CAG-3’ and 5’-GTC TCG TGG GCT CGG AGA TGT GTA TAA GAG ACA G-3’ to primers 19F and 407R, respectively) (Amplicon et al., 2013). PCR was performed under the following conditions: initial denaturation at 95 °C for 1 min; 30 cycles of 95 °C for 15 s, 51 °C for 15 s and 72 °C for 30 s; and final extension at 72 °C for 6 min. PCR products were purified using the HighPrep PCR Clean-up System (MagBio Genomics, Gaithersburg, USA) and used for library preparation with the Nextera XT kit (Illumina, San Diego, USA). The samples were sequenced in the MiSeq platform using the MiSeq Reagent Kit v3 (Illumina).

### *nifH* amplicon sequence analysis

The *nifH* sequences obtained in the previous step were analyzed with the QIIME 2 pipeline version 2023.5 (Bolyen et al., 2019). Primers and adapter sequences were trimmed from the raw reads with the q2-cutadapt plugin. The results were denoised and representative amplicon sequence variants (ASVs) were selected with q2-dada2. The representative ASVs were compared against the *nifH*, *chlL* and *bchX* database included with NifMAP v. 1.2 (Angel et al., 2018) using FrameBot 1.2.0 (Cole et al., 2014), and those that were closer to *nifH* than the other genes were retrieved with a custom script. A reference *nifH* database was prepared from the nifHdada2 dataset v. 2.0.5 (Heller et al., 2014; Moynihan & Furbo Reeder, 2023) and the representative ASVs were taxonomically identified using the q2-feature-classifier plugin. The ASVs identified as potentially belonging to plastids or mitochondria as well as homologs of *nifH* (e.g., *bchX* and *chlL*) were filtered from the dataset. The ASV sequences were aligned with MAFFT (Katoh & Standley, 2013) and a phylogenetic tree was reconstructed by the maximum likelihood method with FastTree (Price et al., 2010) using the q2-phylogeny plugin. Alpha and beta diversity analyses were performed with the q2-diversity plugin.

### Quantitative PCR of the *nifH* gene

Standards were produced by amplifying the *nifH* gene from the bacterium *Ensifer meliloti* (Dangeard) Young using the modified 19F and 407R primers under the conditions described above. The PCR products were visualized, pooled and then cleaned using the QIAquick PCR Purification Kit (Qiagen). qPCR was performed with the Brilliant III Ultra-Fast SYBR Green qPCR Master Mix (Agilent Technologies, Cedar Creek, USA) using the modified 19F and 407 primers. All reactions were performed in three technical replicates on 96-well plates, which also included serial dilutions of the standard in two technical replicates, with concentrations decreasing exponentially from 1.35·10^9^ and 1.35·10^3^ copies·µL^−1^ using the LightCycler 96 Real-Time PCR System (Roche Diagnostics, Mannheim, Germany). Reaction conditions were as follows: initial denaturing at 95 °C for 10 min; 40 cycles of 95 °C for 30 s, 51 °C for 30 s, and 72 °C for 30 s; and a final cycle of 95 °C for 1 min, 55 °C for 15 s and 95 °C for 15 s. For each sample, the procedure was repeated until a quantification cycle variance of less than 1 was achieved between technical replicates (Bustin et al. 2009), after which the average of the three corresponding concentrations was calculated.

### Adjustment of *nifH* quantifications

The sequencing results showed that the Illumina-adapted 19F and 407R primers were not highly specific to *nifH*, also capturing the paralogous *chlL*/*bchL* photosynthesis gene, especially in moss chloroplasts. Therefore, the *nifH* qPCR average concentration values were corrected by multiplying absolute values by the fraction of *nifH* reads recovered from each sample as estimated in the sequence analyses (Angel et al., 2018). For samples identified in the order *Nostocales*, the corrected values were divided by two to avoid overestimations, since many members of this order have two distinct *nifH* copies (Chen et al., 2022). Finally, the number of *nifH* copies for specific taxa was estimated by multiplying the qPCR results for each sample by corresponding relative abundances obtained with the *nifH* sequence analyses.

### Statistical analyses and visualization

The data from the acetylene reduction assays, *nifH* sequencing and qPCR were statistically analyzed with R 4.2.2 ((https://www.R-project.org/) in Rstudio 2022.12.0 (http://www.rstudio.com/). The normality of the datasets was visualized with Q-Qplots and evaluated with Shapiro-Wilk’s test (Shapiro & Wilk, 1965), and the data was transformed with the Yeo-Johnson method (Yeo & Johnson, 2000) using the package car 3.1.3 (Fox & Weisberg, 2019). Differences between N_2_ fixation rates, ASV numbers or *nifH* copies and different hosts, sites or incubation regimes were evaluated with one-way and two-way analyses of variance (ANOVA) (Fisher, 1918), and significant variation between factors were evaluated by the Tukey’s Honestly Significant Difference post-hoc test (Tukey, 1949). Linear regressions (Galton, 1886) were used to investigate the potential relationships between N_2_ fixation, temperature, moisture and *nifH* copies. The results were visualized with the R packages ggplot2 3.5.1 (Wickham, 2016), cowplot 1.1.1 (https://wilkelab.org/cowplot/) and patchwork 1.2.0 (https://patchwork.data-imaginist.com/). Since differences between acetylene reduction analyses performed at different time points were negligible, the data obtained in different weeks were analyzed together.

## RESULTS

### Effects of site, host species, temperature and moisture on N_2_ fixation

Overall, samples collected in ABK, where feather mosses experience lower precipitation, produced an average of 7.9(±0.7) nmol ethylene·g^−1^·h^−1^ in acetylene reduction assays. The average N_2_ fixation rate in ABK samples less than half of than those under higher annual precipitation in the LTJ and KTJ sites, which presented averages of 16.4(±1.5) and 18.1(±1.3) nmol ethylene·g^−1^·h^−1^, respectively (one-way ANOVA, *p* < 0.001, *F*_2,1346_ = 53.08) (Figure 1A). Considering all samples irrespective of origin, a significant, positive relationship was observed between N_2_ fixation and temperature (linear regression, *p* < 0.001, *r^2^* = 0.16) as well as N_2_ fixation and moisture, albeit with a very low explanatory power (*p* < 0.001, *r^2^* = 0.02) (Figures 1B and 1C). No interaction between host species and site was found, but a significant interaction between temperature and moisture was found when all samples were considered (two-way ANOVA, *p <* 0.001, *F*_1,1345_ = 29.16), thus suggesting highest N_2_ fixation rates at the highest temperature (25 °C) and moisture levels (150 %).

**Figure 1.**
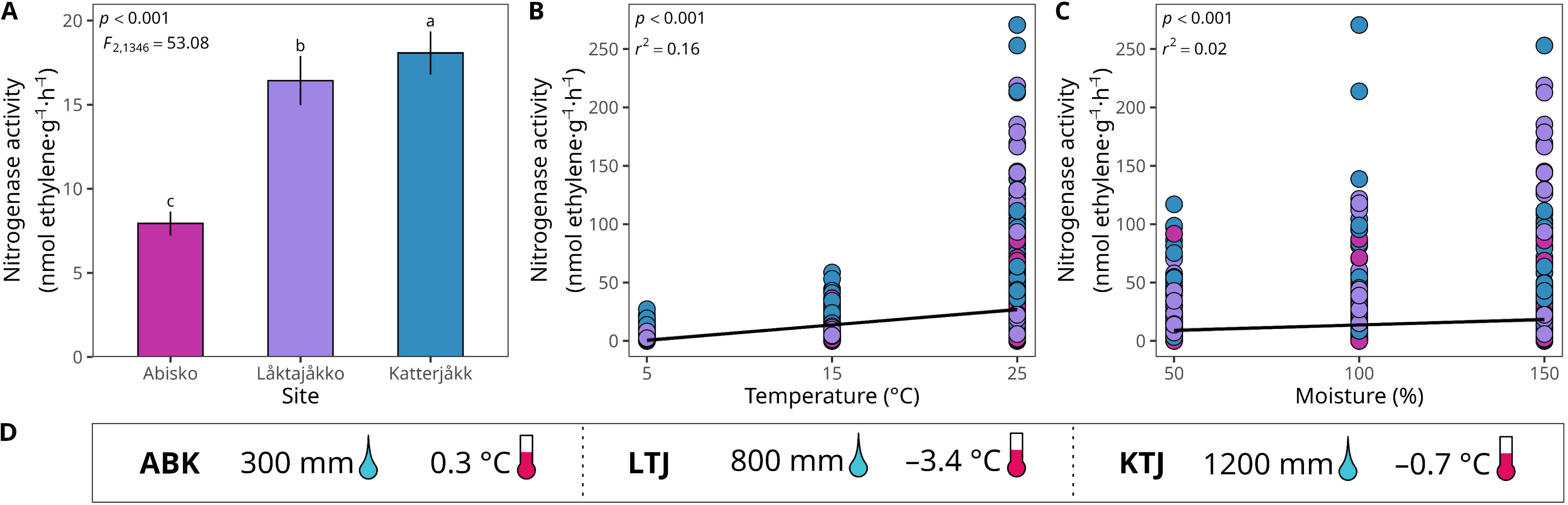
Nitrogenase activity associated with feather moss samples collected across a natural precipitation gradient in northern Sweden as estimated with acetylene reduction assays. A) Moss-associated nitrogenase activity in each site, from driest to wettest. Bars indicate averages, while vertical lines illustrate standard errors. Different letters above bars indicate significant differences in one-way ANOVAs according to Tukey’s Honest Significant Difference post-hoc test. *n* = 540. B) Relationship between increasing temperatures and the nitrogenase activities associated with feather mosses; *n* = 450. C) Relationship between increasing moisture levels and nitrogenase activities; *n* = 450. D) Annual precipitation and temperature averages in each site. ABK: Abisko; LTJ: Låktajåkko; KTJ: Katterjåkk.

When considering the host species individually, we found a trend for both feather moss species from all sites to increase nitrogenase activity with increasing temperature (Supplementary Figure 1). However, unlike communities associated with *P. schreberi*, the N_2_-fixing activities of microbes in association with *H. splendens* were not affected by increasing moisture levels (Supplementary Figure 2). Nevertheless, a significant interaction between temperature and moisture was observed in multiple linear regressions for both host species (Figure 2). *H. splendens* had higher N_2_ fixation rates than *P. schreberi* at 5 °C (one-way ANOVA, *p* < 0.001, *F*_1,448_ = 102.73) and 15 °C (*p* < 0.001, *F*_1,448_ = 41.11), while the opposite was observed at 25 °C (Figure 3A-C). Similarly, *H. splendens* presented higher nitrogenase activity at 50 % (*p* < 0.001, *F*_1,448_ = 87.58) and 100 % moisture (*p* < 0.001, *F*_1,447_ = 18.97), while *P. schreberi* had higher N_2_ fixation rates at 150 % moisture (Figure 3D-F).

**Figure 2.**
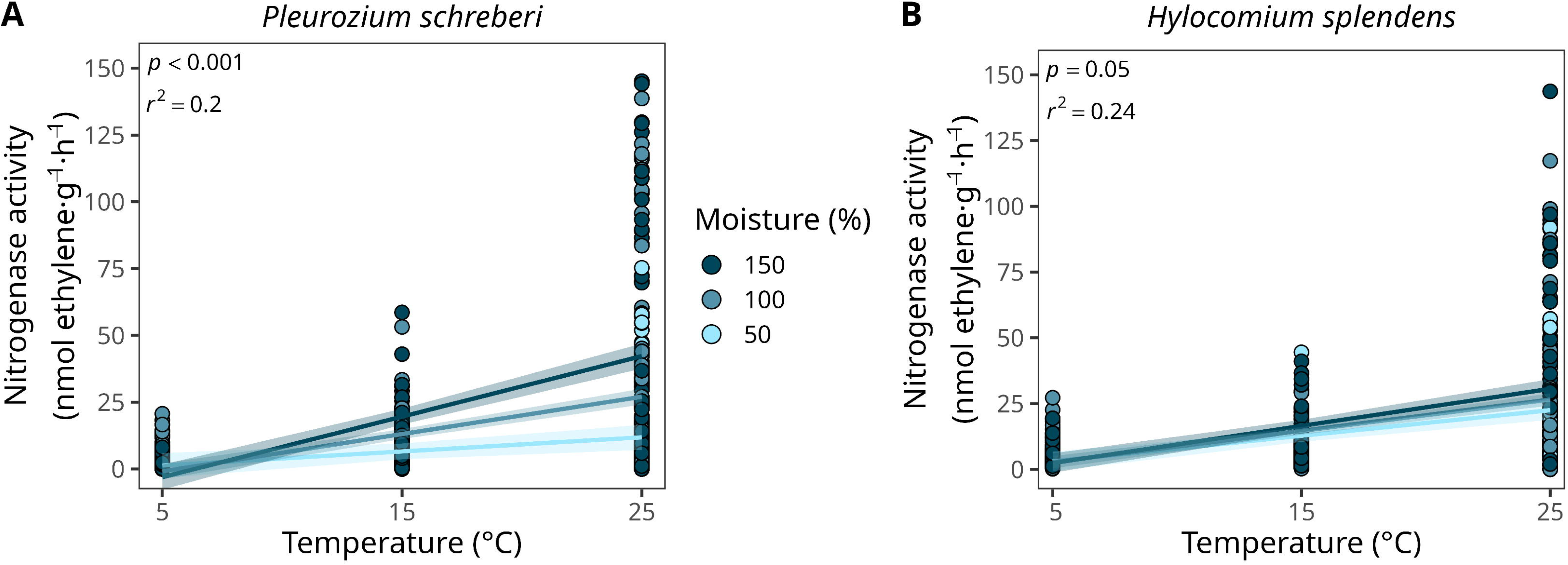
Relationship between nitrogenase activity and temperature by moisture levels in subarctic moss-associated microbial communities based on multiple linear regression analyses. (A) Nitrogenase activities associated with the feather moss *P. schreberi*; *n* = 270. B) Nitrogenase activities in association with *H. splendens*; *n* = 180.

**Figure 3.**
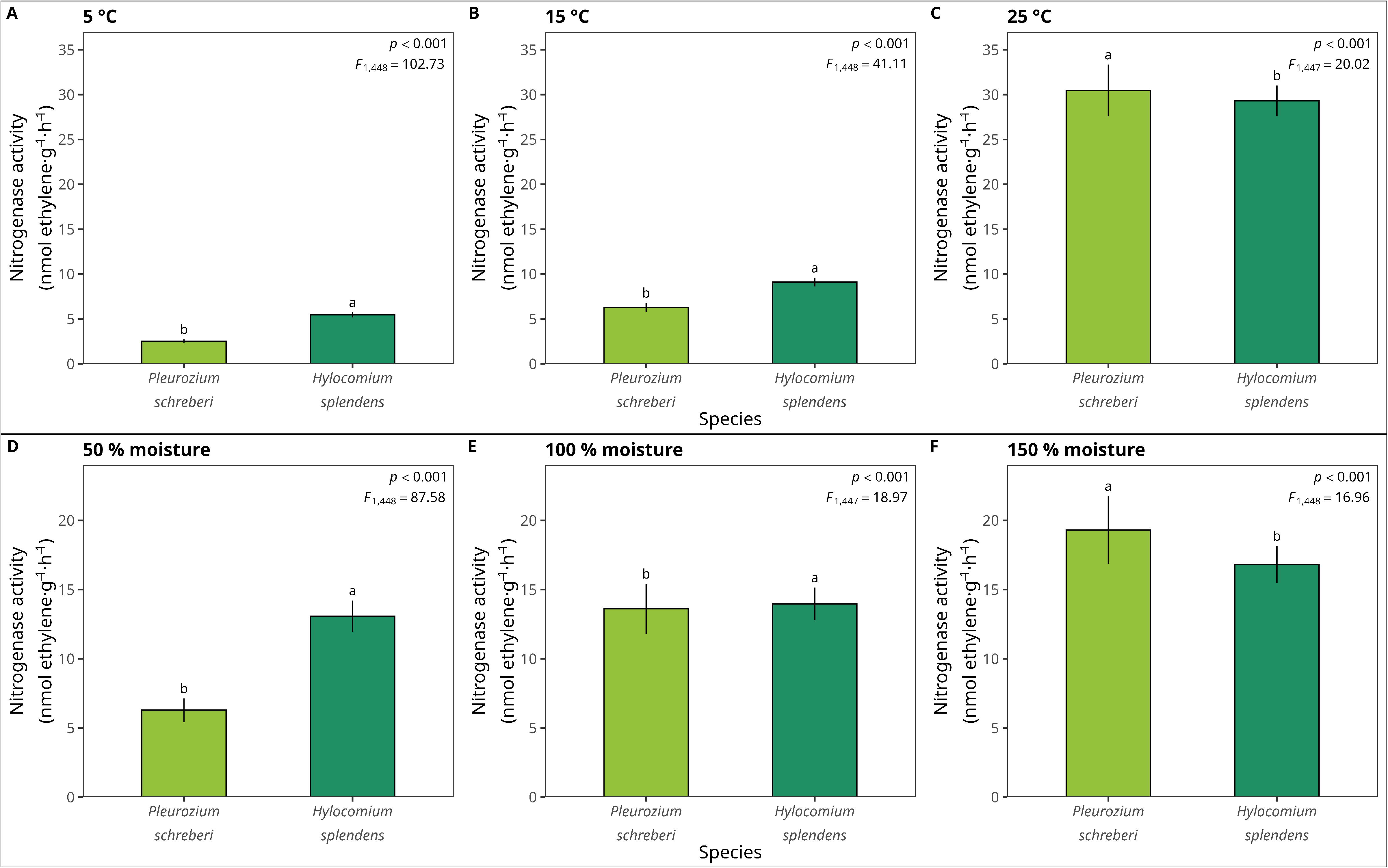
Nitrogenase activity associated with the feather moss species *P. schreberi* and *H. splendens* after 8 weeks of incubation under temperatures of 5 (A), 15 (B) and 25 °C (C) or under 50 (D), 100 (E) or 150 % moisture (F). Averages and standard errors are represented as bars and vertical lines, respectively, while significant differences in one-way ANOVA tests according to Tukey’s HSD tests are indicated by different letters. *n* = 180 and 270 for *H. splendens* and *P. schreberi*, respectively.

Site of origin was also a factor in how *P. schreberi* responded to temperature: samples from the site with the lowest precipitation, in ABK, showed only a slight increase in N_2_ fixation activity with increasing temperatures (linear regression, *p* = 0.001, *r^2^*= 0.04), while the samples from the other sites and from *H. splendens* showed sharper increases (Supplementary Figure 1). Moisture also had lower influence on N_2_ fixation associated with *P. schreberi* samples collected from ABK than on those from the other sites, and no influence on *H. splendens* samples from any site (Supplementary Figure 2).

### Diversity of N_2_-fixing communities per host and site

Sequencing of the *nifH* gene in he feather moss microbiome samples resulted in a total number of 23 528 140 read pairs. Overall, 701 different representative ASVs were identified after filtering among the different samples, with a mean length of 330 bp (Supplementary Table 1). Alpha diversity analyses based on the *nifH* sequences showed that *H. splendens* samples had a significantly higher diversity for N_2_-fixing microbes than *P. schreberi* samples, considering Shannon entropy index (Kruskal-Wallis, *p* < 0.001, H = 32.38), observed ASVs (*p* < 0.001, H = 45.19), Pielou’s evenness (*p* < 0.001, H = 13.97) and Faith’s phylogenetic diversity (*p* = 0.042, H = 4.14). Samples from the site with the peak annual precipitation rate, in KTJ, had significantly higher Shannon diversity index (*p* < 0.001, H = 21.65) and Pielou’s evenness (*p* < 0.001, H = 33.27) than those collected in both ABK and LTJ. Alpha diversity results are summarized in Table 1 and Supplementary Table 2.

**Table 1.** Statistical differences in the alpha diversity of microbial communities associated with feather mosses along a precipitation gradient in northern Sweden incubated under varying conditions. Statistical differences were assessed using the Kruskal-Wallis test across all treatments. Sigificant differences (*p* < 0.05) are highlighted in bold. Values are rounded up.

| Factor | Group |  |  | Shannon entropy |  | Observed features |  | Pielou's evenness |  | Faith's phylogenetic diversity |  |
| --- | --- | --- | --- | --- | --- | --- | --- | --- | --- | --- | --- |
|  |  |  |  | *p* value | H | *p* value | H | *p* value | H | *p* value | H |
| Host species | *Hylocomium splendens* | *Pleurozium schreberi* | — | **0.001** | 32.38 | **0.001** | 45.19 | **0.001** | 13.97 | **0.042** | 4.14 |
| Annual precipitation (mm) | 300 | 800 | 1200 | **0.001** | 21.65 | **0.001** | 13.99 | **0.001** | 33.27 | **0.035** | 6.76 |
| Temperature (°C) | 5 | 15 | 25 | 0.169 | 3.57 | 0.220 | 3.04 | 0.349 | 10.39 | 0.175 | 3.49 |
| Moisture (%) | 50 | 100 | 150 | 0.161 | 3.66 | 0.215 | 3.08 | **0.017** | 8.17 | **0.003** | 11.14 |

Beta diversity analyses also showed that the N_2_-fixing communities differed between the feather moss species (Figure 4) in a quantitative sense, as per Bray-Curtis dissimilarity (PERMANOVA, *p* = 0.001, pseudo-*F* = 11.99) and weighted UniFrac (*p* = 0.004, pseudo-*F* = 7.04) distances, as well as a qualitative sense, as per the Jaccard (*p* = 0.001, pseudo-*F* = 14.41) and unweighted UniFrac (*p* = 0.001, pseudo-*F* = 7.99) distance tests. In addition, significant differences between each of the three communities in the ABK, LTJ and KTJ sites were observed in Bray-Curtis dissimilarity (*p* = 0.001, pseudo-*F* = 19.92), Jaccard distance (*p* = 0.001, pseudo-*F* = 9.93), unweighted UniFrac (*p* = 0.001, pseudo-*F* = 4.86) and weighted UniFrac (*p* = 0.002, pseudo-*F* = 7.13) distance tests. Results for PERMANOVA analyses of the beta diversity between all groups are summarized in Table 2.

**Figure 4.**
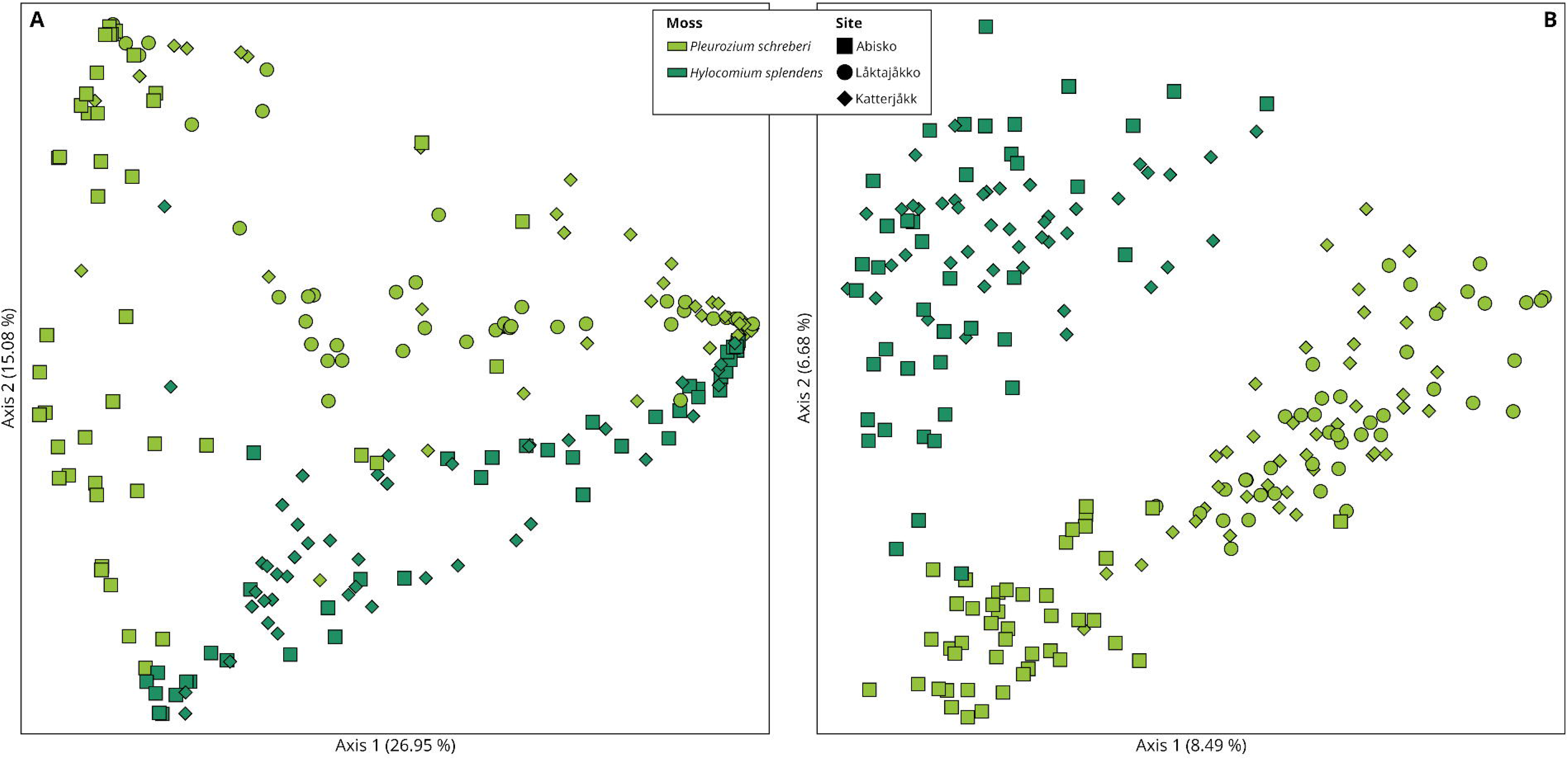
Principal component analyses of beta diversity analyses of the N_2_-fixing communities associated with feather mosses across a precipitation gradient in northern Sweden. A) Bray-Curtis dissimilarities illustrating the dissimilarities in ASV abundance between samples. *n* = 74. B) Jaccard distances depicting how community composition differs between samples, based on ASV presence or absence. *n* = 74.

**Table 2.** Statistical differences in the beta diversity of microbial communities associated with feather mosses along a precipitation gradient in a Swedish subarctic tundra. The analyses were performed with PERMANOVA tests considering all groups. Significant differences (*p* < 0.05) are highlighted in bold. Values are rounded up.

| Factor | Group |  |  | Bray-Curtis dissimilarity |  | Jaccard distance |  | Weighted UniFrac |  | Unweighted UniFrac |  |
| --- | --- | --- | --- | --- | --- | --- | --- | --- | --- | --- | --- |
|  |  |  |  | *p* value | Pseudo-*F* | *p* value | Pseudo-*F* | *p* value | Pseudo-*F* | *p* value | Pseudo-*F* |
| Host species | *Hylocomium splendens* | *Pleurozium schreberi* | — | **0.001** | 11.99 | **0.001** | 14.41 | **0.004** | 7.04 | **0.001** | 7.99 |
| Annual precipitation (mm) | 300 | 800 | 1200 | **0.001** | 19.92 | **0.001** | 9.93 | **0.002** | 7.13 | **0.001** | 4.86 |
| Temperature (°C) | 5 | 15 | 25 | **0.001** | 9.76 | **0.001** | 4.12 | **0.001** | 17.83 | **0.001** | 5.34 |
| Moisture (%) | 50 | 100 | 150 | 0.394 | 1.02 | **0.018** | 1.48 | 0.201 | 1.56 | **0.001** | 3.56 |

### Differences in diversity among incubation regimes

Based on the *nifH* analyses, incubations under different temperatures did not lead to significant changes in the alpha diversity of the moss-associated N_2_-fixing communities (Table 1). However, dissimilarities between the communities in all temperature treatments were observed in beta diversity analyses in both qualitative (*p* = 0.001, pseudo-*F* = 4.12 and *p* = 0.001, pseudo-*F* = 5.34 for Jaccard distance and unweighted UniFrac distances, respectively) and quantitative (*p* < 0.001, pseudo-*F* = 9.76 and *p* = 0.001, pseudo-*F* = 17.83 for Bray-Curtis dissimilarity and weighted UniFrac distances) metrics (Table 2, Supplementary Table 3). Similarly, different moisture treatments did not result in significant differences in Shannon enthropy and observed ASVs, although evenness and phylogenetic diversity did significantly decrease in the 150 % moisture samples (*p* = 0.017, H *=* 8.17 and *p* = 0.003, H *=* 11.14, respectively) (Table 1). Dissimilarities in N_2_-fixing communities exposed to different moisture levels were revealed by beta diversity analyses in qualitative terms only, as indicated by Jaccard distance (*p* = 0.001, pseudo-*F* = 2.47 and *p* = 0.006, pseudo-*F* = 1.66, respectively) and unweighted UniFrac (*p* = 0.001, pseudo-*F* = 4.82 and *p* = 0.002, pseudo-*F* = 4.60) distances (Table 2).

### Differences in N_2_-fixing community composition

Sequencing of the *nifH* gene showed that, overall, cyanobacteria were the dominant members of the N_2_-fixing communities evaluated, with an average relative abundance of 62.2(±1.7) % across all samples (Figure 5). The second most abundant group of N_2_ fixers in the samples were unclassified bacteria (those not presenting any significant similarity with bacterial references in the *nifH* database used), representing an average of 27.9(±1.6) % of the overall community (Figure 5).

**Figure 5.**
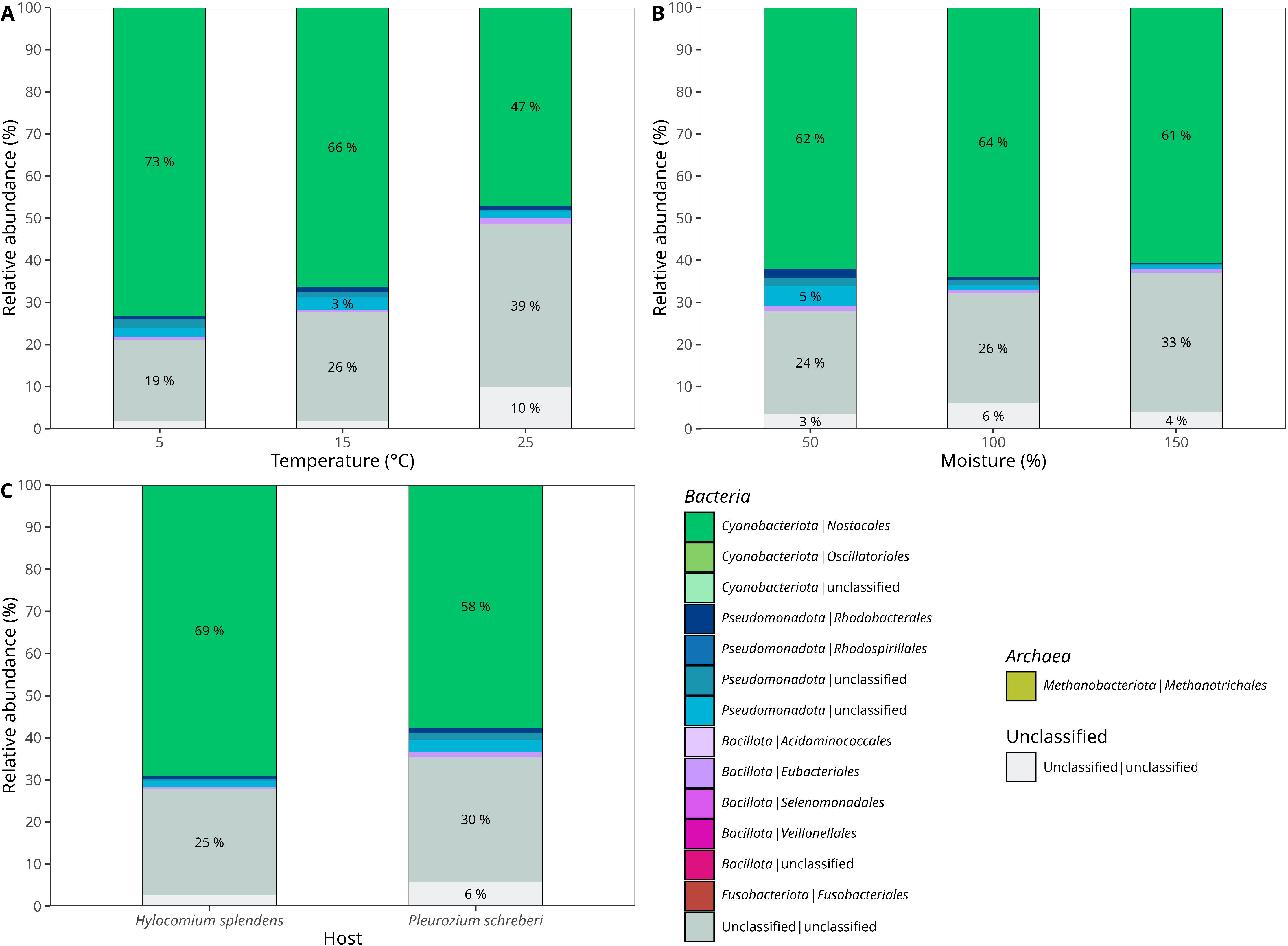
Relative abundances of ASVs for N_2_-fixing bacteria and archaea associated with subarctic feather mosses in relation to different temperatures (A), moistures (B) or hosts (C) based on *nifH* gene analyses. *n* = 74. ASVs for which no significant similarity at the taxonomic level of order or above was found are noted as “unclassified” at their lowest level identified. Sequences related to eukaryotic organisms were removed before plotting.

The relative abundance of cyanobacterial amplicon sequence variants (ASVs) differed significantly between *H. splendens*, with an average of 69.1(±2.7) %, and *P. schreberi*, with 57.7(±2.9) % (one-way ANOVA, *p* < 0.001, *F*_1,220_ = 11.48). These differences were not observed between the different sites across the precipitation gradient, neither between incubations under different moisture levels. However, incubations at 25 °C did lead to a significant decrease in the relative abundance of cyanobacteria (47.0±2.4 %) in comparison with incubations at 15 (66.4±2.8%) and 5 (73.2±2.8 %) °C (one-way ANOVA, *p* < 0.001, *F*_2,219_ = 27.77). On the other hand, the temperature of 25 °C increased significantly the relative abundance of unclassified bacteria (38.6±2.6 %) in comparison with samples incubated at 15 °C (25.9±2.7 %) and 5 °C (19.2±2.5 %) (*p* < 0.001, *F*_1,219_ = 14.88). While there was no difference between the relative abundance of unclassified bacteria in the different hosts, the site with the lowest annual precipitation rates (ABK) also had a significant lower relative abundance of these bacteria (22.2±2.6 % against 33.2±2.5 and 30.8±2.4 in KTJ and LTJ, respectively) (*p* = 0.004, *F*_1,219_ = 5.61).

### *nifH* gene quantification

As expected, there was a positive relationship between the number of copies of the *nifH* gene in the different samples as measured by qPCR and their overall N_2_ fixation rates observed in acetylene reduction assays, although the analysis had low explanatory power (linear regression, *p* < 0.001, *r^2^*= 0.06) (Figure 6A). The number of *nifH* gene copies in the samples originating in different sites along the precipitation gradient also followed a similar pattern as the one observed for N_2_ fixation, but only in association with *P. schreberi* shoots (Figure 6B and C)*. P. schreberi* samples from ABK presented an average number of copies (3.2·10^4^) significantly lower than the ones from the LTJ (1.2·10^5^) and KTJ (1.8·10^5^) sites, which did not statistically differ from one another (one-way ANOVA, *p* < 0.001, *F*_2,130_ = 8.43) (Figure 7B). The number of *nifH* copies identified as cyanobacteria and unclassified bacteria also increased along with the precipitation gradient for *P. schreberi* samples, while *nifH* copies from unclassified bacteria actually decreased significantly in *H. splendens* communities from the site with the highest annual precipitation, KTJ (Supplementary Figure 3). On the other hand, the temperature of 25 °C tended to decrease the number of cyanobacterial *nifH* copies (Figure 7A and C), while 150 % moisture tended to increase them (Figure 7B and D) in both host species. The different temperature and moisture treatments did not significantly affect the number of *nifH* gene copies for unclassified bacteria.

**Figure 6.**
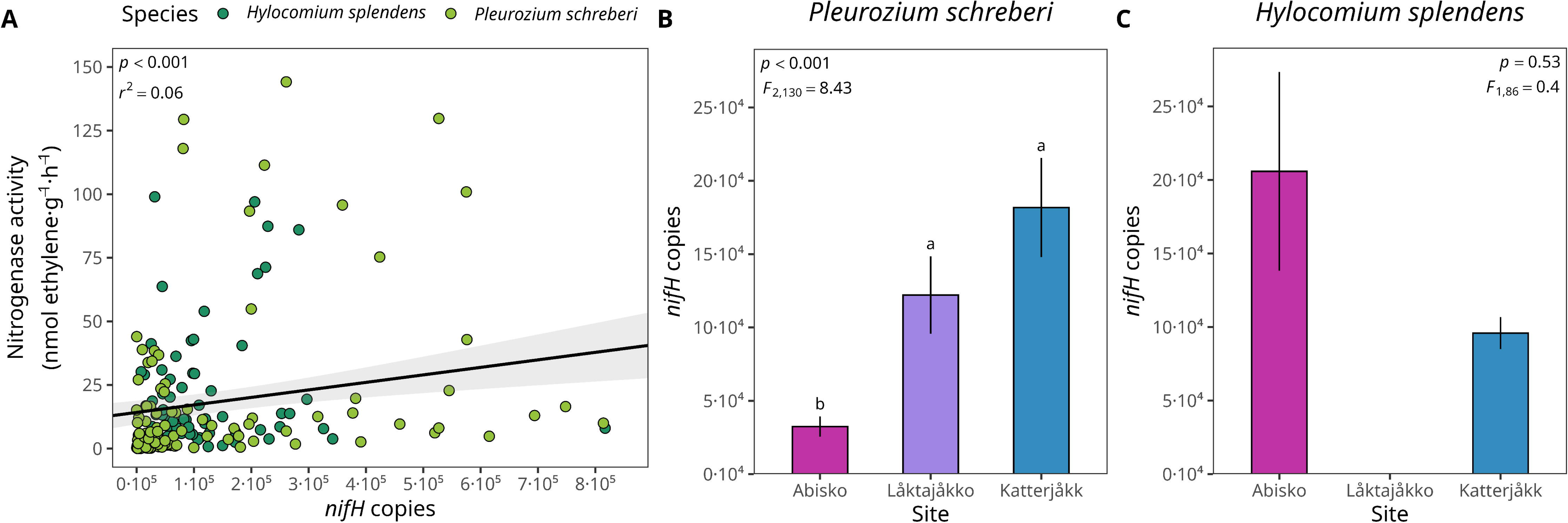
Copies of the nitrogenase gene *nifH* in microbial communities associated with feather mosses from Swedish subarctic tundra as quantified by qPCR. A) Relationship between *nifH* copies and nitrogenase activity based on a linear regression analysis; *n* = 225. B) Number of *nifH* copies expressed in microbial communities associated with *P. schreberi* from three sites forming a precipitation gradient; *n* = 45. C) Differences in the number of copies of the *nifH* gene expressed in communities associated with *H. splendens* samples collected in sites with different annual precipitation rates; *n* = 45.

**Figure 7.**
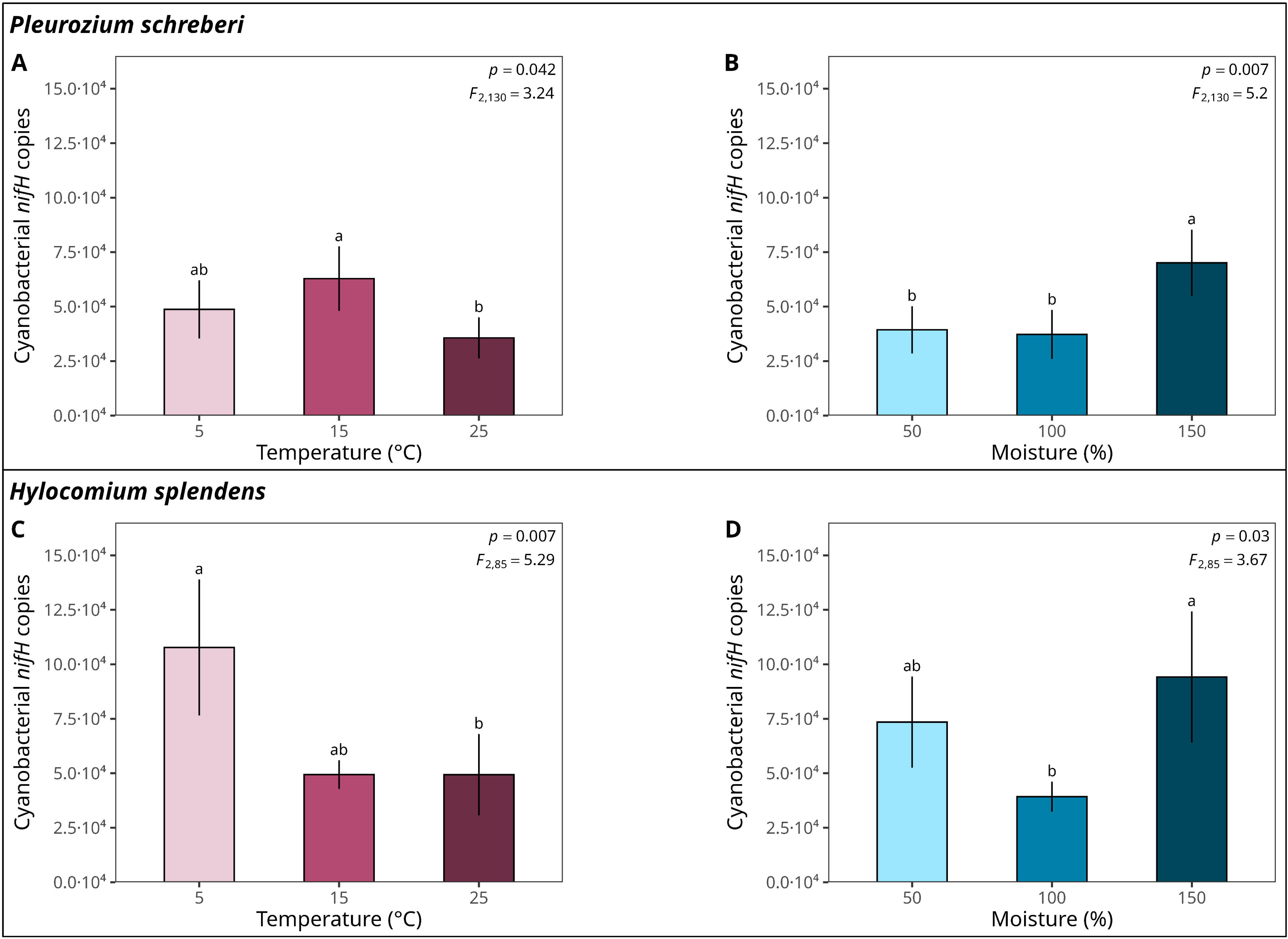
Number of *nifH* gene copies for N_2_-fixing cyanobacteria associated with feather mosses from northern Sweden. A) Effects of different temperatures on the expression of the *nifH* gene by cyanobacteria associated with *P. schreberi*. B) Number of cyanobacteria *nifH* gene copies expressed in association with *P. schreberi* under different moisture levels. C) Number of cyanobacterial *nifH* gene copies expressed in association with *H. splendens* incubated under different temperatures. D) Variation in the number of copies for the *nifH* gene by N_2_-fixing cyanobacteria associated with *H. splendens* under different moistures. Bars and vertical lines represent averages and standard errors, respectively. Treatments receiving different letters were significantly different in one-way ANOVA tests according to Tukey’s HSD post-hoc analyses. *n* = 45.

## DISCUSSION

Our first hypothesis, which anticipated that mosses collected from sites with higher annual precipitation in the Swedish subarctic would exhibit higher N_2_ fixation rates, was supported (Figure 1A), despite their lower mean annual temperature. Since moisture was normalized between samples before the experiments, the observed effect was unlikely to be due to a carryover of water content and its physiological effects in the feather mosses from the field conditions. More likely, it was a direct result of the lasting influence of their environments of origin on the microbial communities associated with the feather moss samples.

Furthermore, *nifH* gene copy numbers and nitrogenase activity in *P. schreberi* samples from the driest site, ABK, showed only slight increases in response to higher temperatures and moistures (Supplementary Figures 1 and 2). This suggests that the legacy of extended exposure to field factors continue to influence the N_2_-fixing communities associated with *P. schreberi* even when conditions change. In contrast, the *H. splendens*-associated communities from different sites showed consistently higher nitrogenase activity with greater relative abundance of cyanobacterial ASVs and little variation in *nifH* gene copies, suggesting more stable communities that only require a lower level of moisture for their activities, not benefiting from extra water. In addition, alpha diversity was highest in association with mosses from the site with the highest annual precipitation (KTJ). This suggests that, overall, precipitation favors the diversity and N_2_-fixing activities of feather moss microbiomes in the subarctic.

Environmental conditions have often been shown to directly impact the microbiomes of vascular plants, creating long-lasting effects on the composition and function of their microbial communities (Trivedi et al., 2022). Our results indicate that environmental factors also have profound effects on the microbial communities associated with non-vascular plants. Considering that mosses display highly plastic phenotypes that respond strongly to environmental fluctuations (Mohanasundaram & Pandey, 2022) and that plants and their microbiomes function as a holobiont (Vandenkoornhuyse et al., 2015), it is not surprising that moss microbiomes are also significantly impacted by environmental factors. Furthermore, compared to vascular plants, mosses provide fewer colonization sites for microbes, which in most cases are restricted to living epiphytically on their hosts (Alvarenga & Rousk, 2022). This may further expose moss microbial communities to environmental pressures. Our findings also suggest that as climate change alters temperature and precipitation patterns in the subarctic, it will probably leave imprints on moss microbiomes and their ecological functions.

Overall, our second hypothesis, which stated that the N_2_ fixation rates associated with the feather moss samples would be positively influenced by temperature and moisture, was confirmed (Figures 1, 2 and 3). However, there were noticeable differences between the communities associated with the two moss species. Sites with higher moisture seem to support a greater abundance of N_2_-fixing microbes in *P. schreberi*, especially cyanobacteria and unclassified bacteria (Figure 7). In *H. splendens* samples, increasing moisture levels did not lead to higher nitrogenase activity (Supplementary Figure 2D and 2E). This was also reflected in the consistent microbial composition across treatments in this species. This is counterintuitive, but could be partially explained by the differences in the microbial communities between feather moss species (Figure 4), suggesting that *H. splendens* may host more resilient N_2_-fixing microbes than *P. schreberi*.

Not much is known about the microbial diversity associated with mosses across different ecosystems. Nevertheless, considerable differences between the composition of bacterial communities in the microbiomes of different host species have been observed, possibly resulting from co-diversification or ecological filtering driven by moss traits (Holland-Moritz et al., 2018; 2021). Plant species and genotypes are directly related with the different effects microbiomes and their hosts have on each other (van Rensburg et al., 2024), potentially influencing microbial responses to the environment in unique ways. Therefore, despite occupying similar niches in the subarctic tundra and boreal forests that sometimes even lead the the two moss species to physically grow together, these mosses have significantly distinct communities that react differently to their environments.

Temperature had a clear impact on both nitrogenase activity and diazotrophic community composition. Nitrogen fixation increased with temperature across both moss species, but *H. splendens* consistently exhibited higher nitrogenase activity at 5 °C and 15 °C under 50 % moisture. At 25 °C, nitrogenase activity increased in the communities of both moss species, particularly in *P. schreberi* (Figure 3), despite reductions in the relative abundance of cyanobacteria (Figure 5). Beta diversity analyses showed shifts in microbial community, possibly resulting from unclassified bacteria becoming more prevalent, particularly in *P. schreberi*. Moisture also had more pronounced effects in *P. schreberi* communities, which significantly increased nitrogenase activity at 150 % moisture, likely resulting from higher numbers of cyanobacterial ASVs and *nifH* gene copies. In contrast, *H. splendens* was more resistant to changes in moisture, with nitrogenase activity, community composition and *nifH* gene copies remaining relatively stable across moisture levels. These findings suggest that while *P. schreberi* is more responsive to moisture fluctuations, *H. splendens* maintains a more resilient nitrogen-fixing community under varying moisture conditions. Finally, as also predicted by our third hypothesis, we found a significant relationship between temperature and moisture levels in relation to cyanobacterial ASVs (Figure 6). In general, the effects of temperature on the N_2_-fixing activities of cyanobacteria seems to be dependent on their environment and habitat. For example, N_2_-fixing cyanobacteria from aquatic environments tend to grow and perform better in temperatures of 25 °C or a bit higher (Thomas & Litchman, 2016), while the optimal temperature for the N_2_-fixing activities of cyanobacteria in boreal forests may vary between 16 and 27 °C (Rousk et al., 2017b). Some works have found that arctic and subarctic cyanobacteria in various habitats have a comparable temperature optimum for N_2_ fixation around 25 and 32 °C (Zielke et al., 2002; Salazar et al., 2022), while another observed higher N_2_-fixing activities in temperatures under 14 °C (Rousk et al., 2018). Our results provide another evidence that some cyanobacteria inhabiting tundra ecosystems may actually fix more N_2_ under colder temperatures.

Virtually all the N fixed in moss-cyanobacteria symbioses can be retained within host tissues in the short term, but precipitation can cause it to significantly leach to the ground, depending on the host species (Song et al., 2024). This process is known to introduce a significant amount of N into arctic and subarctic soils that can be taken up by other organisms, allowing them to bypass the N limitations that are common in these ecosystems. Based on the results we obtained, however, climate change could decrease the abundance of cyanobacterial ASVs and *nifH* copies in these communities (Figure 5) as it leads to not only enhanced shrub growth outcompeting mosses (Elmendorf et al. 2012), but also to a warmer and more drought-prone Arctic.

This may seem paradoxical, as we did observe that, overall, higher temperatures tend to lead to increased nitrogenase activity (Figures 1, 2 and 3, Supplementary Figure 1). This result was apparently driven by the rise in the abundance of ASVs belonging to N_2_-fixing bacteria from currently unclassified taxa (Figure 5), which was also observed by Holland-Moritz et al. (2018). Similarly, Klarenberg et al. (2022) have observed the partial replacement of N_2_-fixing cyanobacteria with other diazotrophs in the microbiome of the subarctic moss *Racomitrium lanuginosum*, although followed by a decrease in the number of overall *nifH* gene copies in the community. We observed stable *nifH* copy numbers and nitrogenase activity from unclassified bacteria across temperatures despite the change in taxa composition and a drop in cyanobacterial *nifH* gene copies, suggesting that unclassified bacterial taxa may take over the N_2_-fixing role from cyanobacteria without compromising overall nitrogenase activity, thus allowing these associations to remain important players in the biogeochemical cycle of N in a changing Arctic. Driven by increased temperature, this replacement rate may accelerate in the near future due to an increase in the frequency and intensity of hot weather events, as witnessed by the 38 °C recently measured in the Arctic region (World Meteorological Organization, 2021), which is well above the optimal temperatures for N_2_ fixation ever estimated for cyanobacteria from any habitat. On the other hand, this replacement effect seems to be host-dependent, with *H. splendens* showing relatively more stable N_2_-fixing communities that are nonetheless more sensitive to temperature in comparison to those in association with *P. schreberi*.

## CONCLUSIONS

As climate change increases temperatures and the frequency of drought events in the Arctic, the contribution of moss-associated cyanobacteria to N_2_ fixation in the tundra will likely decrease due to its susceptibility to low precipitation. Even though different moss species harbor distinct NLJ-fixing communities, they are all affected by climate change, although not in the same manner. Our findings suggest that currently unclassified bacteria may take over the role of N_2_ fixation from cyanobacteria in feather moss microbiomes, potentially compensating for the decline in cyanobacterial activity. While this shift could help maintain or even enhance N_2_ fixation rates in these communities, the broader ecological consequences of this microbial turnover remain uncertain.

## ACKNOWLEDGEMENTS

The analyses in this work have been performed using the Danish National Life Science Supercomputing Center (Computerome) and the Danish eInfrastructure Cooperation (DeiC) high performance computing resources. We would like to thank Pia A. Pedersen for providing the plasmids used for producing the qPCR standards. We are also grateful to Michael Westbury, Julian Perez and Jakob Russel for their valuable suggestions during the early stages of this project.

## FUNDING

This work was funded by the European Research Council’s Horizon 2020 research and innovation program (grant #947719 to KR), with further support from the Independent Research Fund Denmark (IRFD; #6108-00089). The Danish National Research Foundation supported activities within the Center for Volatile Interactions (VOLT, #DNRF168). The analyses in this work were also funded by the Danish e-Infrastructure Consortium (grants #DeiC-AAU-N1-2024087 and #DeiC-KU-N3-2024088 to DOA).

## SUPPLEMENTARY MATERIAL

**Supplementary Figure 1.**
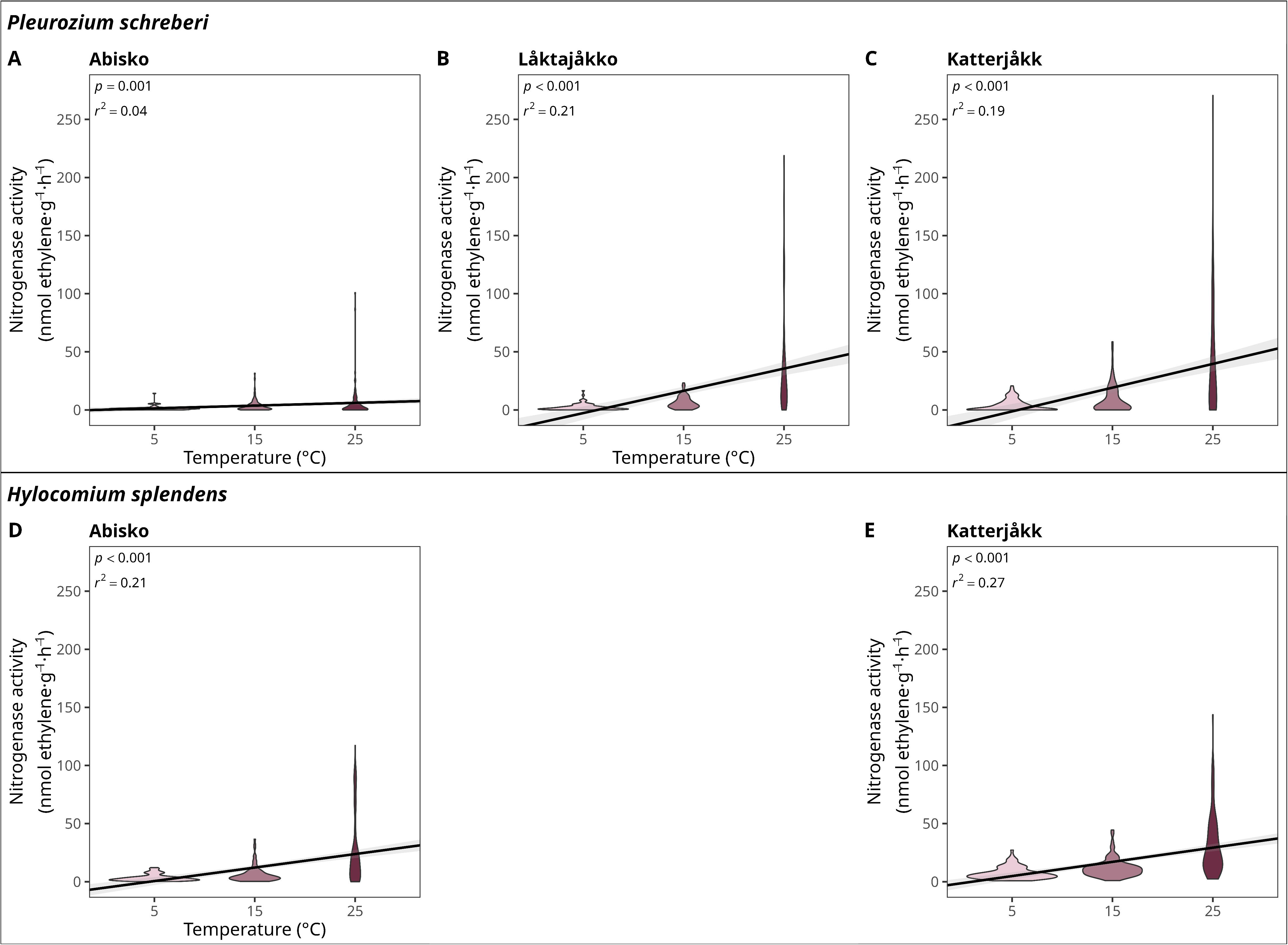
Relationship between temperature and nitrogenase activity associated with samples of the feather moss species *P. schreberi* (A, B and C) and *H. splendens* (D and E) collected in Abisko, Låktajåakko and Katterjäkk, northern Sweden. *n* = 90.

**Supplementary Figure 2.**
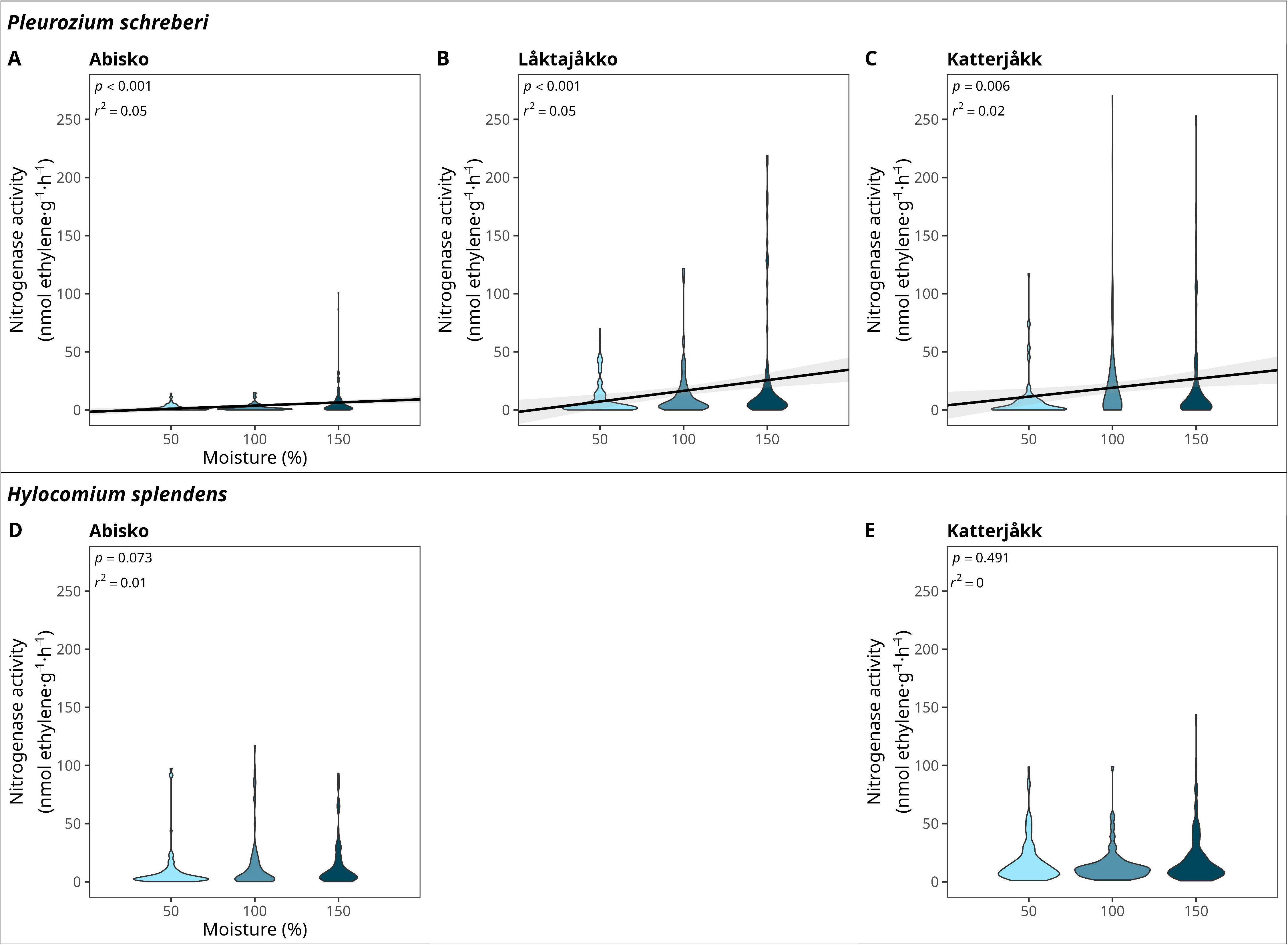
Relationship between moisture and nitrogenase activity in the feather mosses *P. schreberi* (A, B and C) and *H. splendens* (D and E) sampled from a precipitation gradient composed of sites located in the northern Sweden cities Abisko, Låktajåakko and Katterjäkk. *n* = 90.

**Supplementary Figure 3.**
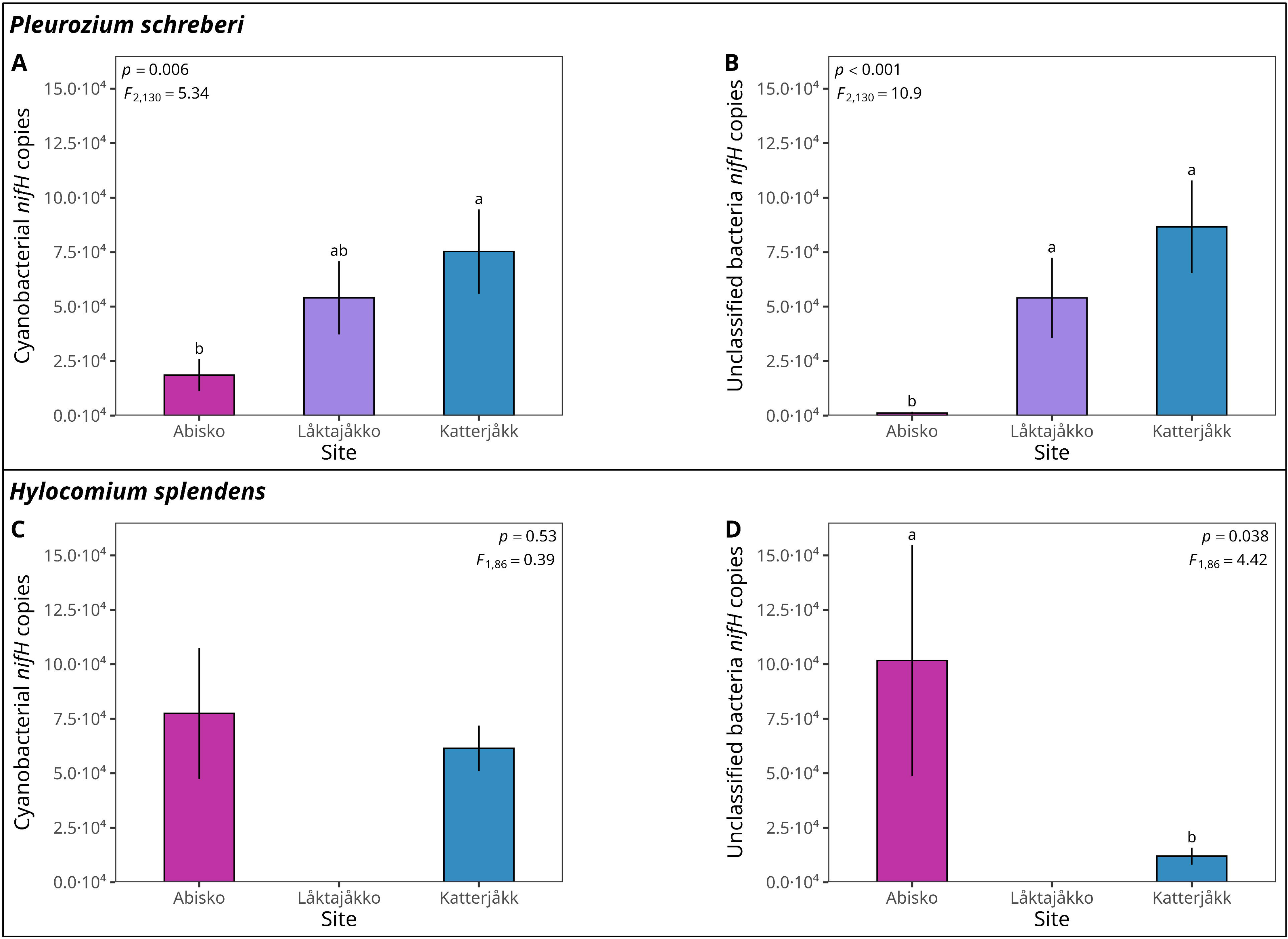
Average number of *nifH* gene copies identified as cyanobacteria or unclassified bacteria associated with the subarctic feather mosses *P. schreberi* (A and B) and *H. splendens* (C and D) along a precipitation gradient in northern Sweden. Bars represent averages, while vertical lines indicate standard errors. Letters above bars indicate significant differences between treatments in one-way ANOVA analyses as per Tukey’s HSD post-hoc test. *n* = 45.

**Supplementary Table 1.** Representative amplicon sequence variants and their taxonomic identification for *nifH* genes in microbial communities associated with subarctic feather mosses in northern Sweden.

**Supplementary Table 2.** Pairwise differences in the alpha diversity of microbial communities associated with feather mosses from a precipitation gradient under different temperature and moisture conditions. Provided are the *p* values obtained in Kruskal-Wallis tests. Significant differences (*p* < 0.05) are highlighted in bold.

**Supplementary Table 3.** Pairwise differences in the beta diversity of microbial communities associated with feather mosses from a precipitation gradient under different temperature and moisture conditions. Provided are *p* values from PERMANOVA tests. Significant differences (*p* < 0.05) are highlighted in bold.

## Notes

### Competing Interest Statement

The authors have declared no competing interest.

